# Neuronal excitability is permanently altered by activity manipulation during an embryonic critical period in *Drosophila*

**DOI:** 10.64898/2025.12.01.691172

**Authors:** Bramwell Coulson, Jacob J. Davies, Tom Pettini, Matthias Landgraf, Richard A. Baines

## Abstract

Neuronal intrinsic excitability provides the baseline that homeostatic mechanisms act to preserve, yet the processes that establish a baseline remain poorly defined. Developmental critical periods (CPs) are thought to play a central role, but the link between early activity and long-term intrinsic properties is not well characterised. To address this, we used the genetic tractability of the *Drosophila* larval locomotor circuit to manipulate individual neurons during an embryonic CP. Following optogenetic excitation or inhibition, during the CP, we assessed intrinsic excitability of the same neurons in third-instar larvae (i.e. ∼5 days thereafter). We compared an excitatory premotor interneuron (A27h), an inhibitory premotor interneuron (A31k), and a motor neuron (aCC). Both interneurons exhibited anti-homeostatic responses: excitatory perturbation increased intrinsic excitability, while inhibitory perturbation decreased it, effects that persisted throughout larval development. In contrast, motor neurons showed no significant changes under the same conditions, revealing cell type-specific sensitivity to early activity. These findings build on the general principles of the functional relationships between CP activity and neuronal excitability and how intrinsic excitability is not passively set but actively shaped during these windows, with long-lasting, neuron-specific consequences. More broadly, our results highlight how developmental perturbations can alter the excitatory–inhibitory balance of mature neural circuits that may contribute to the aetiology of neurodevelopmental disorders.

## Summary

Individual neurons set, and thereafter maintain, an appropriate level of excitability to ensure network stability. Indeed, without such an ability (often referred to as neuronal homeostasis), the self-reinforcing properties of Hebbian-based changes would likely culminate in network destabilisation (Lee and Kirkwood, 2019; Cooper and Bear 2012; Turrigiano et al., 1994; Turrigiano, 1999). Many key questions remain to be answered including when, and how, do neurons encode their intrinsic excitability?

Accumulating evidence implicates, but does not directly show, that intrinsic neuronal excitability is specified during neural development, with activity perturbations during developmental critical periods (CPs) being sufficient to permanently alter neuronal excitability (Zeng et al., 2021; Lybrand et al., 2021; Marguet et al., 2015; Giachello and Baines, 2015; Hunter et al., 2024). For example, optogenetic reduction of network activity, during an embryonic CP in the *Drosophila* embryo, can overcome the hyperexcitability induced by a gain-of-function Na_V_ mutation that normally results in a predisposition to electroshock-induced seizure in mature larvae (Giachello and Baines; 2015). Activity manipulation either before, or after this defined CP is without effect. Measurement of network properties suggest that hyperexcitability during this same CP is sufficient to permanently alter the excitation-inhibition balance of the mature neural circuit, favouring excitation (Hunter et al., 2024). These observations support the hypothesis that neurons track activity they experience during a CP and use this ‘measure’ to set an appropriate level of excitability that then becomes ‘locked-in’ when the CP closes. We show in this study that optogenetic manipulation of individual and identified interneurons, during a defined developmental CP, permanently alters their intrinsic excitability. By contrast, optogenetic activity manipulation after the CP has closed is without effect. Thus, we provide direct experimental evidence to validate this hypothesis.

## Results

Capitalising on the *Drosophila* connectome project, our previous work identified mono-synaptically-connected premotor interneurons and their motoneuron target (Giachello et al., 2022). In this study, we selectively manipulate activity, using optogenetics, of two of these interneurons: a cholinergic excitor (A27h) and a GABAergic inhibitor (A31K), along with their common motoneuron target (aCC). Initially, optogenetic manipulation is restricted to a defined and well-characterized 2 h CP that occurs between 17 -19 h after egg laying (see Giachello and Baines, 2015 for a complete description of this CP) (Figure 1A). The effect of this manipulation is measured ∼5 days later at the late larval wandering third instar stage (L3), by recording the membrane excitability of the manipulated neurons using whole-cell patch clamp (Figure 1B – E).

**Figure 1:**
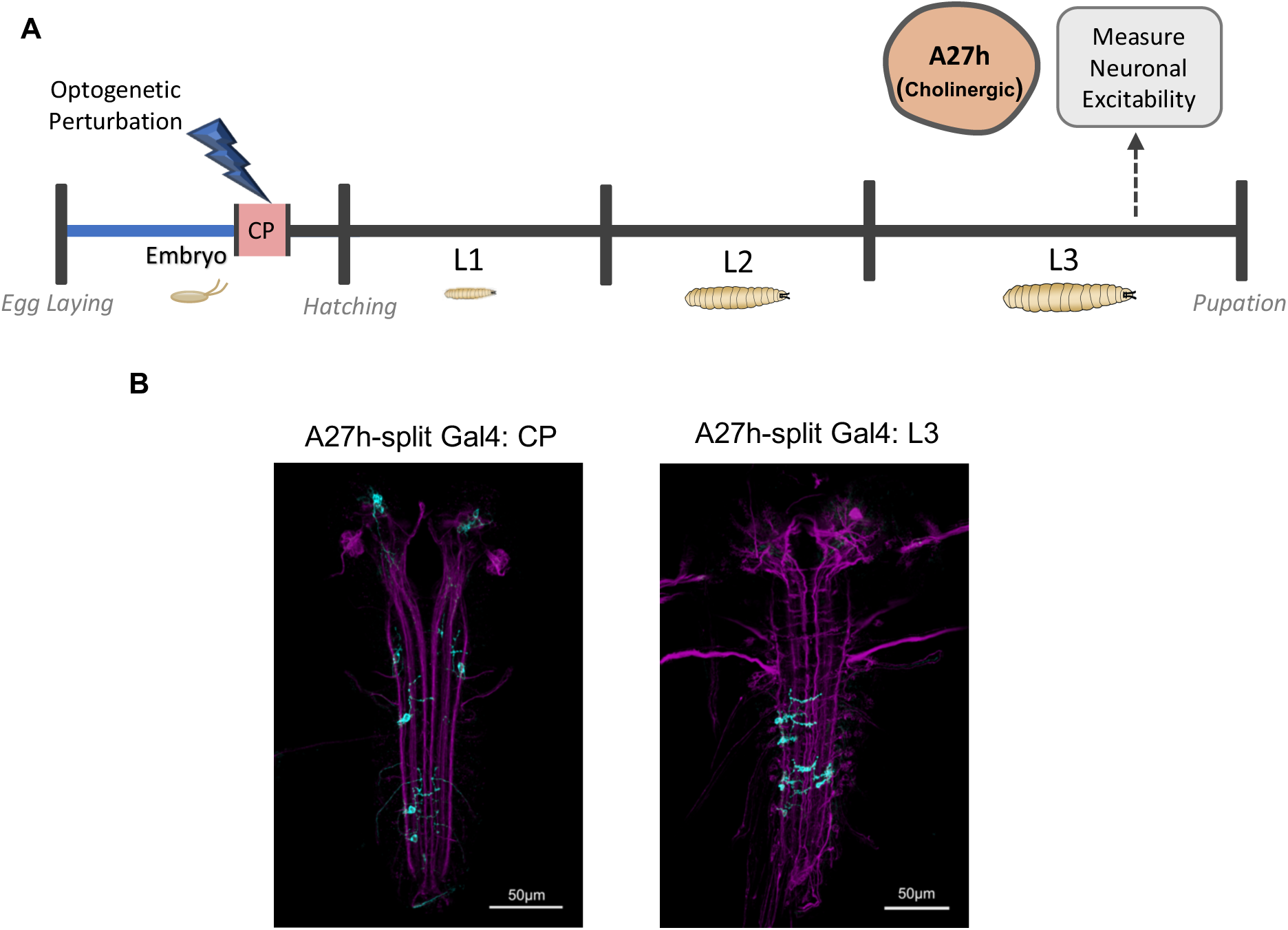
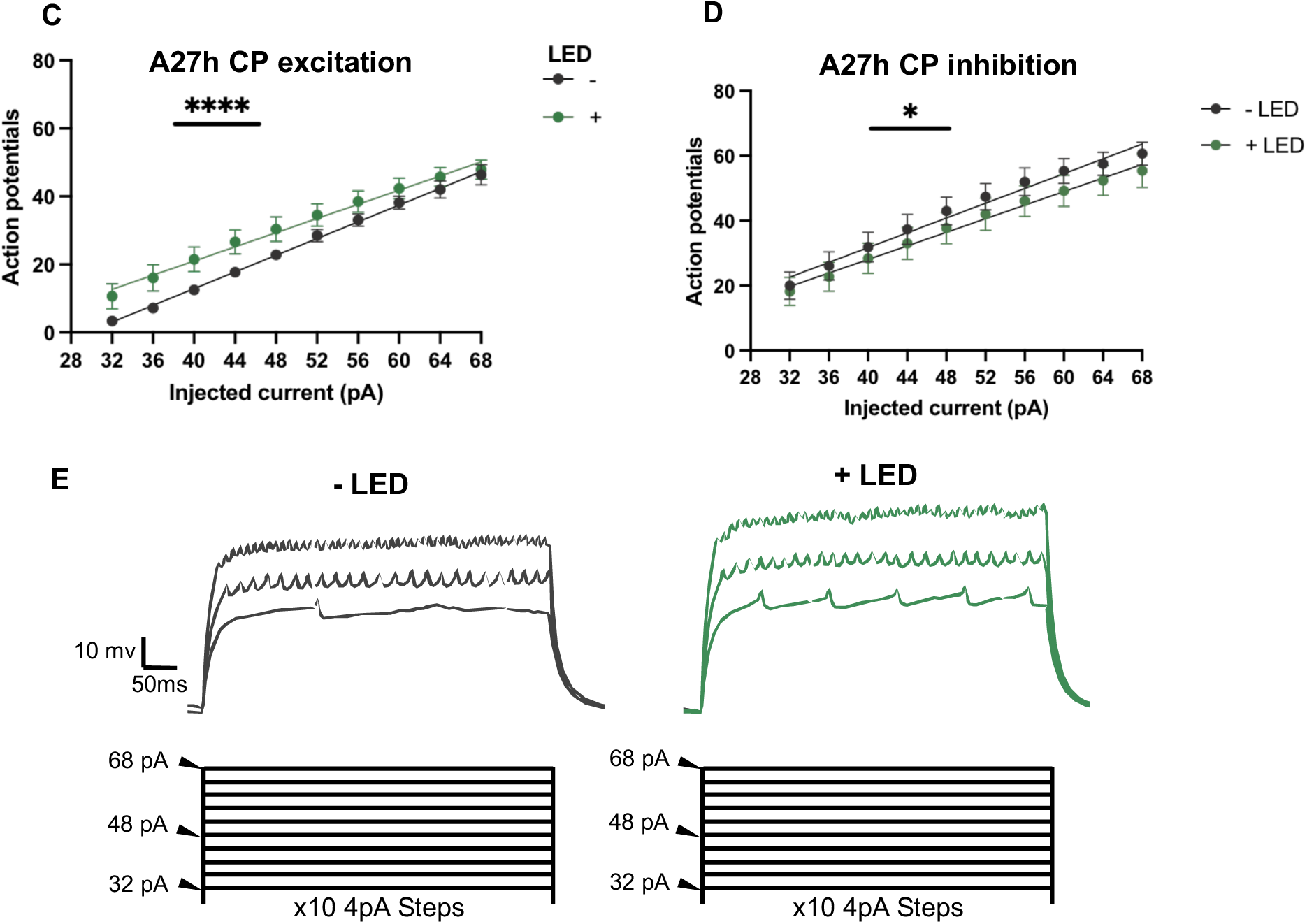
Optogenetic manipulation of the A27h cholinergic excitatory interneuron between 17-19 h AEL bi-directionally alters neuronal excitability. **A**: A27h interneurons were either optogenetically excited or inhibited during the embryonic CP. A27h neurons were then patched at L3 to compare excitability to non-manipulated controls. **B**: Gal4/UAS expression in A27h interneurons at 19 h AEL (left, during the CP), and 72 h AEL (right, at L3) in the ventral nerve cord (VNC). **C**: Optogenetic excitation between 17-19 h AEL increases intrinsic excitability of A27h in L3 larva (-LED: slope 1.229, R^2^ 0.92, n = 6, +LED: slope 1.04, R^2^ 0.64, n = 8, comparison of intercepts, (F(1, 139) = 24.22, p < 0.0001)). **D**: Optogenetic inhibition between 17-19 h AEL decreases intrinsic excitability of A27h in L3 larva, (-LED: slope 1.14, R^2^ 0.55, n = 9, +LED: slope 1.05, R^2^ 0.44, n = 9, comparison of intercepts, (F(1, 177) = 5.73, p = 0.01)). **E**: Representative traces comparing A27h excitability across three current steps (32 pA, 48 pA, and 68 pA) after excitation during the CP (+LED, green), compared to non-manipulated controls (-LED, black).

A 2 h optogenetic excitation of the excitatory interneuron A27h, that coincides with the embryonic CP (17 -19 h AEL, Figure 1A-B), results in increased membrane excitability in these same neurons when recorded some 5 days later at L3 (Figures 1C, E and S1). By contrast, optogenetic inhibition of this interneuron during the CP results in reduced membrane excitability at L3 (Figures 1D, E and S1). Analysis of key cell parameters; cell capacitance (a measure of cell size), input resistance (a measure of membrane integrity) and resting membrane potential (indicator of neuronal ‘health’) show no significant differences, indicative of no gross changes to cell size or viability (Figure S1 B-C).

We repeated the same 2 h embryonic CP experimental manipulation on the GABAergic inhibitory A31k (Figure 2A and B). We observed the same qualitative effects. Thus, optogenetic excitation during the embryonic CP resulted in increased membrane excitability of A31k at L3 (Figure 2C and S2). By contrast, optogenetic inhibition was sufficient to reduce membrane excitability (Figure 2D and S2). Again, key cell parameters remained constant (Figure S2 B-C).

**Figure 2:**
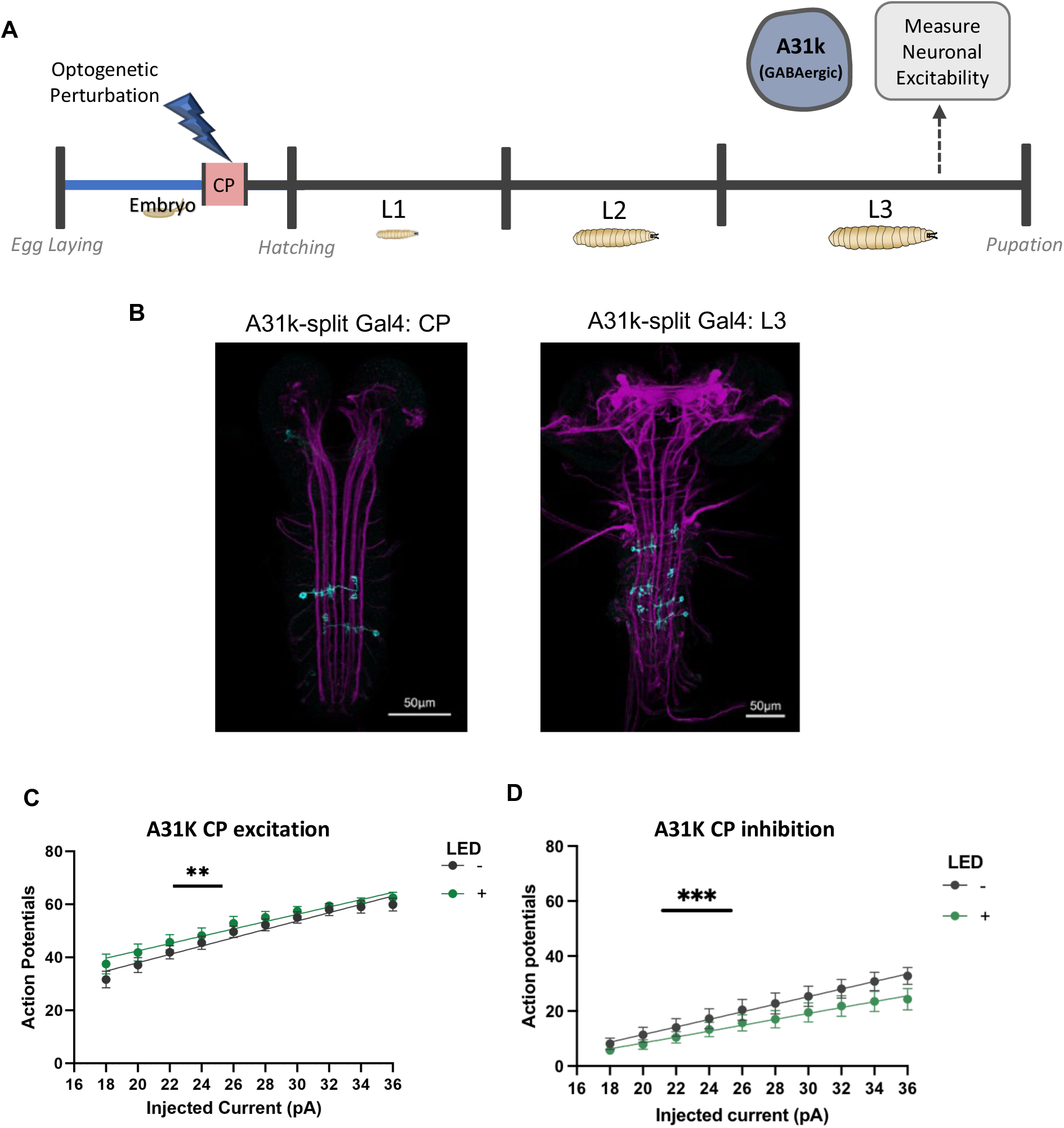
Optogenetic manipulation of the inhibitory GABAergic A31k interneuron between 17-19 h AEL bi-directionally alters neuronal excitability. **A**: A31k interneurons were either optogenetically excited or inhibited during the embryonic CP. A31k neurons were then patched at L3 to compare excitability to non-manipulated controls. **B**: Gal4/UAS expression in A31k interneuron at 19 h AEL (left), and 72 hr AEL (right) in the VNC. **C**: Optogenetic excitation between 17-19 h AEL increases intrinsic excitability of A31k in L3 larva (-LED: slope 1.38, R^2^ 0.51, n = 10, +LED: slope 1.57, R^2^ 0.58, n = 10, comparison of intercepts, (F(1, 197) = 7.99, p = 0.005)). **D**: Optogenetic inhibition between 17-19 h AEL decreases intrinsic excitability of A31k in L3 larva (-LED: slope 1.38, R^2^ 0.42, n = 9, + LED: slope 1.07, R^2^ 0.32, n = 10, comparison of intercepts, (F(1, 187) = 14.66, p = 0.0002)).

Our previous observations show that activity manipulation either side of the embryonic CP is without significant effect to locomotor network stability indicative that, for effects to be permanent, they must occur whilst the CP is ‘open’ (Giachello and Baines, 2015). Thus, we repeated optogenetic manipulation of A31K (GABAergic inhibitor) at 20 -22 h AEL, a time when our previous work shows that this CP has closed (Figure 3A). As expected, manipulation of A31K, at this later stage of embryogenesis, failed to affect membrane excitability of this neuron when recorded at L3 (Figure 3B and S3).

**Figure 3:**
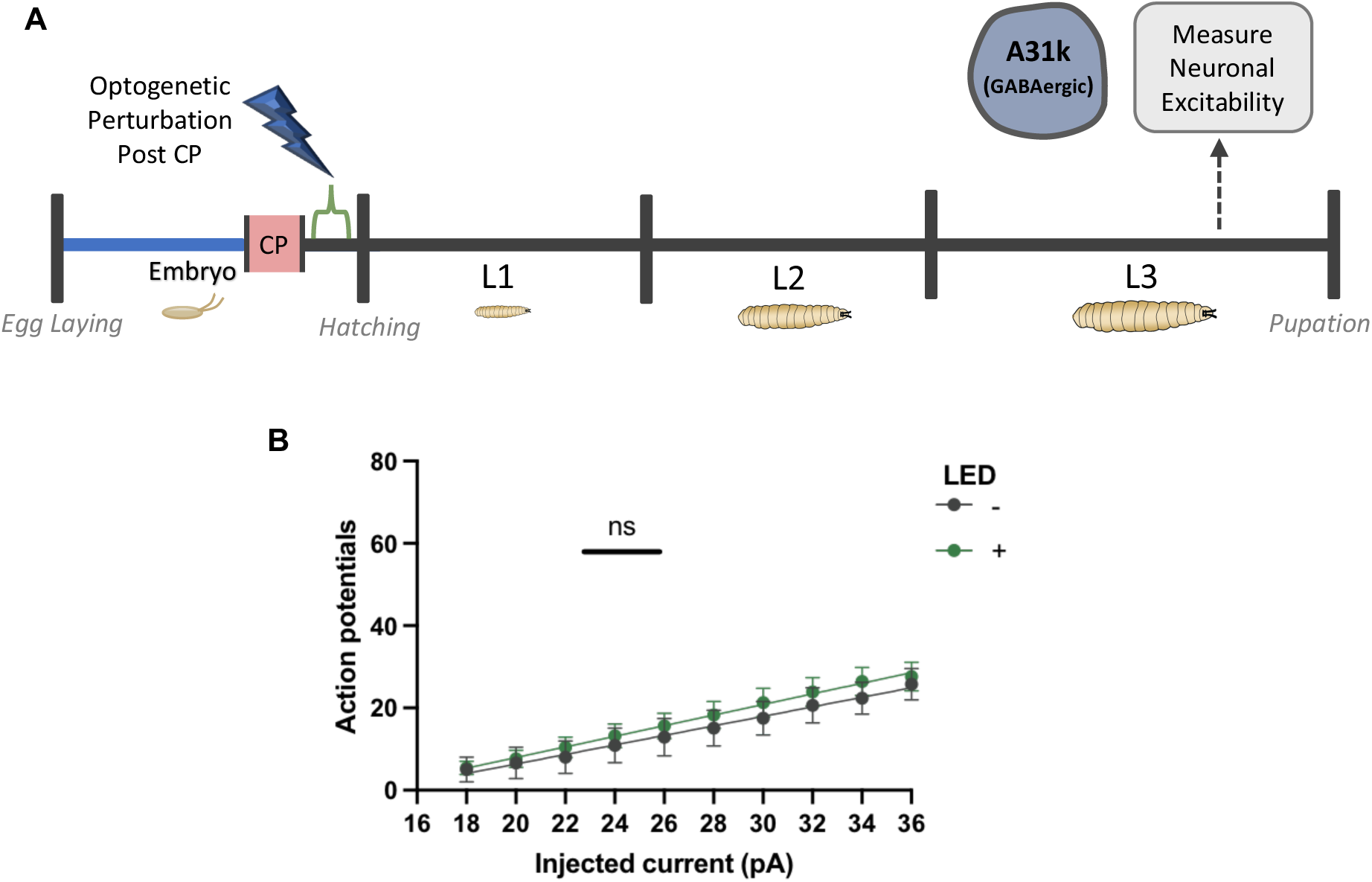
Optogenetic manipulation post CP does not alter intrinsic excitability. A: A31k interneurons were optogenetically inhibited during the embryonic CP. A31k was then patched at L3 to compare excitability to non-manipulated controls. B: Optogenetic inhibition of A31k between 19-21 h AEL does not alter intrinsic excitability in L3 larva (-LED: slope 1.16, R^2^ 0.28, n = 8, +LED: slope 0.16, R^2^ 0.4, n = 10, comparison of intercepts, (F(1, 177) = 2.84, p = 0.09)).

### Embryonic CP activity manipulation does not alter motoneuron excitability

Our results above support the hypothesis that developing interneurons track and encode the activity they are exposed to during a CP to, possibly permanently, set their respective intrinsic excitability. To ask whether other neuron types might behave similarly, we activity-manipulated an identified motoneuron (and synaptic target of these two interneurons) termed ’aCC’ (Figure. S4). Notably, optogenetic activity manipulation (either excitatory or inhibitory), during the embryonic CP, did not alter the excitable properties of the aCC motoneuron when recorded at L3 (Figure S4 and S5).

We next investigated how manipulated interneurons might alter their membrane excitability. In previous work, we have shown that membrane excitability of *Drosophila* neurons can be homeostatically regulated through change of the voltage-gated Na^+^ (I_Na_) current expressed by individual neurons (Baines et al., 2001; Baines, 2003; Muraro et al., 2008; Driscoll et al., 2013). The I_Na_ expressed by neurons in *Drosophila* exhibits at least two components: a fast transient (I_NaT_) and a slower persistent (I_NaP_) current component. The voltage step we used for our analysis was optimised to evoke the maximal amplitude of each component, respectively (Lin et al., 2012). We repeated the optogenetic manipulation (both excitation and inhibition) of activity of A31K during the embryonic CP and measured both I_NaT_ and I_NaP_ expressed in this neuron at L3 (Figure 4A). Due to the small cell size of A31k neurons, I_NaP_ amplitude was found too small to measure reliably; subsequently, I_NaP_ was not compared between conditions. For I_NaT_, the effects we observed were mixed: I_NaT_ was reduced in amplitude following optogenetic inhibition, whereas no change was observed to I_NaT_ following optogenetic excitation during the CP (Fig. 4A-D). No differences were observed for either cell capacitance or input resistance between conditions (Figure S6). Thus, whilst a change to I_NaT_ may underlie reduction in intrinsic excitability, a different mechanism seemingly drives increased excitability.

**Figure 4:**
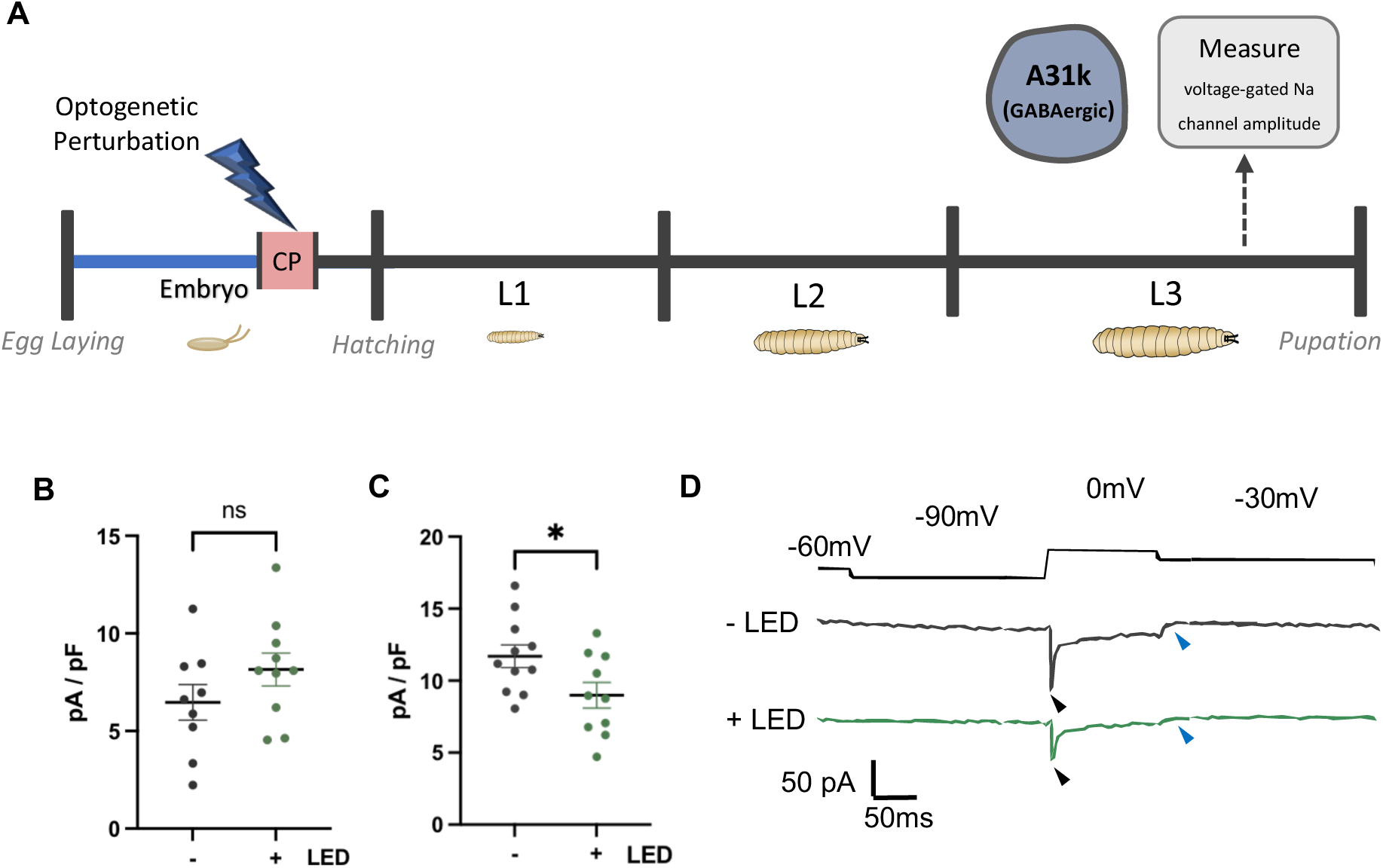
Activity-induced alteration of I_NaT_ may contribute to setting neuronal excitability. **A:** A31k neurons were either optogenetically excited or inhibited during the embryonic CP. The transient sodium current amplitude in A31k was then compared at L3. **B**: Optogenetic excitation between 17-19 h AEL does not alter I_NaT_ current amplitude of A31k in L3 larva (unpaired t test, *t*(17) = 1.365, *p* = 0.193, *n* = 9,10). **C**: Optogenetic inhibition between 17-19 h AEL decreases I_NaT_ in A31k in L3 larva (unpaired t test, *t*(19) = 2.279, *p* = 0.034, *n* = 10,11) **D**: Representative traces comparing A31k I_NaT_ (black arrow head) after CP inhibition (+LED, green) compared to non-manipulated controls (-LED, black). I_NaP_ (blue arrow head) was not compared due to reduced signal-to-noise ratio as a consequence of cell size.

## Discussion

Even though neuronal homeostasis has been extensively studied, key questions remain unanswered. These include ‘when do neurons encode a homeostatic set-point’ and, equally, ‘do all neurons work to a common timeline in this regard’? Our results here, at least as illustrated for two types of interneurons, support the premise that activity exposure during a CP is instrumental for irrevocably setting the functional properties of neurons, including encoding of excitability setpoints, and thus contributing to the excitation:inhibition balance of the mature circuits to which they contribute. Developmental CPs have long been recognised as windows during which the so-called excitation:inhibition balance of mature networks is set, and as a time when activity-manipulation is sufficient to induce permanent change of this property (Hensch, 2005). Although this is well established, the functional relationship between activity during a CP and subsequent effects on neuronal excitability remain unclear.

In this study, we exploited the genetically tractable and well characterised *Drosophila* larval locomotor circuit to activity-manipulate identified neurons during a defined embryonic CP. Using sparsely labelled Split-Gal4 constructs to reduce pleiotropic effects, we showed that transient activity manipulation of individual interneuron types, during a CP, results in permanent changes to the intrinsic excitability of these same neurons as demonstrated by recording 5 days later in mature 3^rd^ instar larvae. Using both excitatory and inhibitory optogenetic manipulations, we report bi-directional changes in intrinsic excitability, with no effect seen when those manipulations were conducted after the CP had closed. These observations are consistent with, and indeed provide evidence for, the specification of excitability set points occurring during a CP.

We note that the A27h cholinergic excitatory interneuron exhibited a greater increase in membrane excitability following optogenetic excitation during the CP, as compared to the change in amplitude of reduced excitability following optogenetic inhibition. The reverse was true for the A31k GABAergic inhibitory interneuron. The difference between how these two neurons respond to CP activity-manipulation is intriguing and may suggest that different neuron types (excitatory vs. inhibitory) respond differentially to CP manipulations. That excitatory interneurons respond preferentially to increased activity during a CP compared to inhibitory interneurons that respond preferentially to reduced activity is seemingly anti-homeostatic. This outcome provides a possible mechanistic understanding for why activity manipulation during a CP may destabilize mature circuits. Indeed, we have previously shown that opposing CP manipulations, regardless of their sign (excitatory or inhibitory), tend lead to similar increases in seizure sensitivity in larval stages (Giachello & Baines, 2015). In those experiments, increased tendency to seizure was used as a proxy for network instability. Likewise, embryonic exposure to either the proconvulsant picrotoxin, or to the anticonvulsant phenytoin, was sufficient to increase larval seizure severity and both treatments showed clear changes to the excitatory:inhibitory balance of the larval locomotor network (Hunter et al., 2024).

The results of this present study strengthen previous findings in which we report that the perturbation of certain neuron types, but not others, during the embryonic CP was sufficient to increase seizure activity (i.e. network instability) in L3 larvae. Specifically, it was shown that optogenetic excitation of either GABAergic or cholinergic neurons was sufficient to induce a marked increase in evoked seizure severity in mature larvae. However, identical optogenetic excitation of motoneurons had no discernible effects (Giachello & Baines, 2015). Our present observations validate this by showing changes in intrinsic excitability in both a GABAergic and a cholinergic interneuron, but no change in the glutamatergic aCC motoneuron. It is tempting to speculate on the mechanistic reasoning for these apparent differences. As the outputs of the locomotor circuit, the role of motoneurons is clearly different to that of interneurons that, collectively, form the central pattern generator (CPG). Thus, to be able to faithfully convey any change of CPG output, it might be expected that motoneurons would be resistant to activity perturbation during a CP. Future investigation will be required to resolve this important issue.

In this study, we explored how activity perturbation alters the intrinsic excitability of individual neurons. Such alteration of activity may reflect a change in the encoding of the so-called homeostatic set point. We report changes in the amplitude of transient sodium current (I_NaT_) in A31k neurons after optogenetic inhibition, but not after excitation, during the CP. This is suggestive of separable mechanisms for hypo- vs. hyper- excitation of this neuron during the CP. A reduction in sodium current amplitude is sufficient to reduce AP firing in response to current injection. Previous work has shown the translational repressor Pumilio (Pum) regulates voltage gated sodium channel expression in *Drosophila* larva motoneurons. Intriguingly, loss of function of *pum* is sufficient to increase one of two separable components of I_Na_ (the persistent but not the transient current component), whereas overexpression of *pum* was shown sufficient to reduce both transient and persistent current (Mee et al., 2004). Thus, whilst we might speculate that activity-mediated change in expression of *pum* might underlie the excitability changes we record in both A27h and A31k, this would need to be tested experimentally.

Intracellular calcium is likely to play a key role in defining a neuronal homeostatic set point and is well-positioned for this potential role due to it being a highly ubiquitous ‘activity-sensing’ signal in the nervous system. The influx of intracellular calcium, mediated by voltage-dependent channels, is crucial for many forms of synaptic and intrinsic plasticity (Lisman et al., 2002; Malenka & Bear, 2004). One hypothesis is that intracellular calcium level may be a proxy for intrinsic excitability (LeMasson et al., 1993; LeMasson et al., 1993; O’Leary et al., 2014). Deviation away from a specific target level of cytosolic Ca^2+^ is sufficient to evoke transcriptional changes that, in turn, tune firing properties (Ibata et al., 2008; Goold & Nicoll, 2010). In keeping with this hypothesis, emerging evidence implicates mitochondrial signalling. Mitochondria are known to regulate neuronal variables, such as synaptic vesicle release and intracellular calcium concentration, with a proposal that mitochondrial Ca^2+^ is maintained at a target level and that change in this level serves as an error signal (Ruggiero et al., 2021).

It remains unclear as to how homeostatic set points are coordinated between single neurons vs. larger neuronal networks (Wen & Turrigiano, 2024). For instance, studies have shown that mean firing rates recover more rapidly at a circuit level than in individual cells following activity perturbation (Slomowitz et al., 2015). Subsequently, measures of intrinsic excitability in individual neurons is only one potential aspect of neuronal homeostasis; activity at the network level also being regulated (Ruggiero et al., 2025). Network regulation appears to have subtle complexities. For example, whilst some neuron types restore their own firing rates following activity perturbation (Hengen et al., 2016), different neuron types seemingly act to restore the firing rates of other neurons (Hengen et al., 2013; Gainey et al., 2018). In the rodent visual cortex, hypo-excitability caused by monocular deprivation is compensated for by the disinhibition of fast-spiking GABAergic neurons (Hengen et al., 2013; Gainey et al., 2018). Probing the interplay between excitability at the single cell vs. the network level may provide important insight into how specific components of neural networks exhibit greater influence on establishing functional activity patterns. Mechanosensory input during early development is important for the normal function of the *Drosophila* larval CPG (Crisp et al., 2011; Zeng et al., 2021). Blocking muscular activity during the stages in which un-patterned muscle activity transitions to patterned activity, results in larvae with altered crawling behaviour, with the transition from un-patterned to patterned locomotor activity being key to the emergence of rhythmic CPG activity (Carreira-Rosario et al., 2021). Likewise, a key pioneer circuit has since been identified that appears to be modulated by proprioception and subsequently highlighting what might be considered a “hub neuron” (the M neuron). Significantly, the M-neuron is electrically coupled to A27h neurons and is involved in integrating proprioceptive activity during early development (Zeng et al., 2021). Collectively, these findings provide some evidence to suggest that different neurons respond differently to CP activity perturbation. Our data add to this perspective, indicating that activity-manipulation during a CP affects interneurons but not motoneurons, and that the magnitude of effect is relative to whether the interneuron is an excitor or an inhibitor.

In summary, we provide experimental evidence to support the hypothesis that some, but certainly not all, developing neurons track the activity they are exposed to during a CP and use this ‘measure’ to specify their intrinsic excitability. Once the CP closes then the same neurons become resistant to change. It is possible that those neurons that follow this developmental program contribute to a greater extent to neural circuit formation and, as such, any imposed changes during a CP are sufficient to disturb the excitation-inhibition balance of the mature circuit.

## Methods

### Drosophila rearing and stocks

Flies were maintained on standard food consisting of (g/L): 7.93 glucose, 7.2 maize, 50 yeast, 8 Agar, 27ml nipagen (10% in EtOH) and 3ml proprionic acid) at 25°C in a 12:12 light/dark cycle. The following lines were used; w; +; GMR94G06-Gal4 (a.k.a. aCC-Gal4), gift from Juan J. Pérez-Moreno (Pérez-Moreno and O’Kane, 2019); +; R20A03-p65ADzp (attP40); R93B07-ZpGDBD (attP2) (a.k.a, A31k-Gal4, gift from Akinao Nose); w[1118]; +; GMR36G02-Gal4 (attP2) (a.k.a. A27h-Gal4); w*; UAS-Chronos::mVenus (attP40); +(a.k.a UAS-Chronos) (BDSC #77115); w*; +; UAS-GtACR1.d-EYFP (attP2) (a.k.a UAS-GtACR1) (BDSC #92983).

### Developmental activity manipulation

4 g of yeast extract powder (Melford, UK) was added to 1950 µl of ddH2O together with 50 µM of 100 mM all-trans retinal (Sigma-Aldrich, UK), a cofactor necessary for activation of channelrhodopsin. Approximately 1 ml of yeast paste was placed onto the surface of a standard apple grape agar plate which was then attached to the laying pot.

Stocks were selected and combined as a 1:2 ratio of Gal4-line males to UAS-line virgin females, with flies being added to an acrylic egg-laying cage using a funnel. The cage was connected to a grape agar plate (containing food and retinal if required) to form a laying pot. Laying pots were left at 18°C for 48 hr to allow flies to acclimate and ingest food (and all-trans retinal). After 48 hr, pots were moved to 25°C and the agar plate changed at 10:00 and 14:00 h every day for 4 successive days (Monday–Thursday). Embryos were discarded at 10:00 and a fresh clutch was collected until 14:00, constituting embryos that were laid during this 4-hr period. For manipulations post CP, embryos were collected during a 1 hr laying window (10:00 -11:00). Embryos were collected from the surface of the apple agar plates using a wet paintbrush >72 h after the laying pot was set up (Tuesday–Thursday) to ensure laid embryos contained all-trans retinal sufficient for optogenetic activation. Following collection, embryos (of either sex) were split into two groups. Each was placed on the centre of a small (5 cm diameter) grape agar plate, ensuring embryos were laid flat and not overlaying. Both agar plates were placed into a cooled incubator (LMS, UK) at 25°C. One agar plate was placed in a plastic container covered in tin foil to ensure no light could penetrate, with these embryos constituting no-manipulated controls. The other agar plate was placed under a 565 nm Thorlabs LED. A wall socket timer (Timeguard) was connected to a GRASS S48 Stimulator which triggered the LED (1.57 mW / cm^2^) to fire for 100 ms at 3 Hz for 6 hours (03:00 – 09:00) which corresponds to the *Drosophila* larval CP of 17 – 19 h AEL +/- 2 h embryos were then removed from the incubator, removed from the agar plate using a wet paint brush and placed into vials containing standard food (recipe stated above). Vials were encased in tin foil to prevent light exposure for 5 days until L3 stage. For post CP manipulation, the above protocol was carried out using a 1 h laying window (10:00-11:00), with LED stimulation occurring 20 -22 h AEL +/- 1 h (06:00 – 09:00).

### Whole cell patch clamp electrophysiology

Third instar (L3) larvae (of either sex) were dissected under saline (135mM NaCl (Fisher Scientific), 5mM KCl (Fisher Scientific), 4mM MgCl2·6H2O (Sigma-Aldrich), 2mM CaCl_2_·2H_2_O (Fisher Scientific), 5mM TES (Sigma-Aldrich), 36mM sucrose (Fisher Scientific), pH 7.15) using light filtered through a red filter. The ventral nerve cord and brain lobes (CNS) was isolated and transferred to a droplet of saline containing 200 µM Mecamylamine (Sigma, UK). This was used to block postsynaptic nACh receptors to synaptically isolate neurons from excitatory inputs. CNS were laid flat (ventral side up for recordings of both aCC and A27h and dorsal side up for A31k) and glued (GLUture Topical Tissue Adhesive; World Precision Instruments USA) to a Sylgard-coated cover slip (1 to 2mm depth of cured SYLGARD Elastomer (Dow-Corning USA) on a 22 x 22mm square coverslip). The preparation was then placed on a glass slide under a microscope (Olympus BX51-WI). CNSs were viewed under a 60x water-immersion lens. To access soma, 1% protease (Streptomyces griseus, Type XIV, Sigma-Aldrich, in external saline) contained within a wide-bore glass pipette (GC100TF-10; Harvard Apparatus UK, approx. 10 µm opening) was applied to abdominal segments, roughly between A5-A2 for both aCC and A31k and A7-A1 for A31k due to being sparsely expressed (Baines and Bate, 1998). This was done to remove overlaying glia to facilitate access to underlying nerve cell soma. Neurons were identified using an m-venus tag expression for experiments using chronos or EYFP tag expression for experiments using GtACR1, this was done briefly using a 565 nm LED.

Recordings were conducted at room temperature (18°C -22°C) using borosilicate glass pipettes (GC100F-10, Harvard Apparatus) that were fire polished to resistances of 8 – 13MΩ for aCC and 15 -30MΩ for both A27h and A31k when filled with intracellular saline (140 mM potassium-D-gluconate (Sigma-Aldrich), 2 mM MgCl_2_·6H_2_O (Sigma-Aldrich), 2 mM EGTA (Sigma-Aldrich), 5 mM KCl (Fisher Scientific), and 20 mM HEPES (Sigma-Aldrich), (pH 7.4). Input resistance was measured in the ‘whole cell’ configuration, and only cells that had an input resistance ≥ 0.5 GΩ were used for experiments in aCC and A27h, whereas a ≥ 1 GΩ threshold was used for experiments in A31k. Cell capacitance and break-in resting membrane potential was also measured for each cell recorded and compared between groups. Data was captured using a Multiclamp 700B amplifier controlled by pCLAMP (version 10.7.0.3), via an analogue-to-digital converter (Digidata 1440A, Molecular Devices). Trace/s were sampled at 20 kHz and filtered online at 10 kHz. Once patched, neurons were brought to a membrane potential of -60 mV using current injection. For both aCC and A27h recording consisted of 20 x 4 pA (500 ms) current steps, including an initial negative step, giving a range of -4 to +72 pA. The same protocol was used for recordings of A31k; however, 2 pA current steps were used, beginning at -2, with the 20^th^ step being 36 pA. Number of spikes fired were counted and plotted against injected current, across a 10 current step range, matched between experiments. Both cell capacitance and input resistance were compared between conditions to ensure that any observed differences in excitability were not due to differences in either cell size or resistance. Resting membrane was also measured upon break in and compared between conditions.

For sodium current recordings, larva were dissected in standard saline as previously stated. The CNS was then glued to the coverslip and the standard saline was replaced by a sodium current isolation saline (100mM NaCl (Fisher Scientific), 6mM KCl (Fisher Scientific), 10mM MgCl2·6H2O (Sigma-Aldrich) 10mM sucrose (Fisher Scientific), 10 mM HEPES (Sigma-Aldrich), 10 mM 4 aminopyridine (Aldrich), 50 mM triethylamine (Sigma-Aldrich), pH 7.4. After using protease to provide access to A31k neurons (as previously stated), neurons were patched using a sodium conductance internal saline (140 mM CsCH3SO3 ((0.91ml Methanesulfonic acid, 99% (thermo scientific), 2.44ml CsOH sol 50% wt (thermo scientific)), 5 mM CsCl (Sigma-Aldrich), 2 mM MgCl2·6H2O (Sigma-Aldrich), 11 mM EGTA (Sigma-Aldrich), 20 mM HEPES (Sigma-Aldrich), pH 7.4). Cell capacitance was measured upon break in followed by a sodium current protocol recorded in voltage clamp to capture transient sodium current. Cells were held at -90 mV (180 s), followed by 0 mv (100 ms), -30 mV (200 ms), and finally -60 mV (100 ms). Due to the low amplitude of the persistent current measured in A31k, it was not consistently detected, subsequently, only reliable measures of the transient current were compared. The protocol was run multiple times for each cell to generate an averaged traces from up to 5 recordings per cell. The transient sodium current amplitude was measured as the difference in current between the neuron just before the transition from -90 mV and the peak in response after the voltage step to 0 mV; these values were then normalised for current density.

### Sample preparation for immunofluorescence

To visualize Gal4 expression in specific neurons, males from A31K-Gal4, A27h-Gal4 and aCC-Gal4 lines were crossed to virgin females from the UAS-GFP reporter. Crosses were set in laying pots with apple juice agar plates and yeast paste, and incubated at 25°C. For visualising Gal4-driven GFP expression at the embryonic critical period, embryos were collected, dechorionated using 50% sodium hypochlorite solution, and selected at the onset of tracheal filling (∼17 hours AEL) for dissection. For visualising Gal4-driven GFP expression in 72 hours ALH larvae, newly hatched 1^st^ instar larvae were selected under blue light by the presence of gut autofluorescence and then aged for 72 hours on apple juice agar plates with an excess of yeast paste at 25°C, before dissection at 72 hours ALH.

### Immunofluorescence staining of nerve cords

Nerve cords were dissected out in 1X Sorensen’s saline, transferred to Poly-L-Lysine coated coverslips, and fixed in 1X Sorensen’s saline + 4% formaldehyde (Polysciences, 10% Ultrapure methanol-free) for 20 minutes at room temperature. Samples were then washed for 1 hour at room temperature with blocking solution (1X Sorensen’s saline + 0.3% Triton-X-100 + 0.25% BSA). Nerve cords were stained with primary antibodies mouse anti Fasciclin II (DSHB 1D4, used at 1:20) and rabbit anti GFP (ChromoTek PABG1, used at 1:600) diluted in blocking solution and incubated for 16 hours at 10°C with gentle agitation. Samples were washed 4x 15 minutes in blocking solution, then incubated with secondary antibodies goat anti rabbit AlexaFluor568 (Invitrogen A-11036, used at 1:1000) and goat anti mouse STARRED (Abberior STRED-1001, used at 1:1000), diluted in blocking solution and incubated for 12 hours at 10°C with gentle agitation. Samples were washed 4x 15 minutes with 1X Sorensen’s, before storage at 4°C and imaging.

### Immunofluorescence imaging

Image stacks were acquired on a Leica Stellaris 5 confocal microscope, using a 40X/0.80 water dipping objective (for 17 hours AEL nerve cords), or a 20X/0.70 multi-immersion objective with oil (for 72 hours ALH nerve cords). AlexaFluor568 was excited using the 561nm laser at 5% power, with an emission window set to 570-610nm. STARRED was excited using the 638nm laser at 4% power, with an emission window set to 650-720nm. Additional imaging settings were as follows: 1024x1024 pixel format, 16 bit depth, bidirectional scanning, 3x line averaging, pinhole of 1AU, and scan frequency 400Hz. Z-stacks were acquired with 1uM z-interval. Image stacks were maximum projected using FIJI software.

### Quantification & Statistical analysis

Statistical analysis was conducted using GraphPad Prism Software (Version 9.5.1). Data used for pairwise comparisons were tested for normal distribution with Shapiro-Wilk Test. Normally distributed data was analysed using students t-test, whereas non-normally distributed data was analysed using Mann-Whitney U test. Linear regression analysis was conducted to compare the relationship between current injection and action potential firing between conditions. Each neuron type was subjected to both optogenetic excitation and inhibition during the CP. Comparisons between conditions were carried out across a 10 current step range, restricted to linear sections of data, with the 10 current step range being matched between both optogenetic excitation and optogenetic inhibition conditions for each neuron type for consistency. A test for linear trend was compared as part of each linear regression which measured whether the intercepts between the groups were significantly different. For all datasets mean and standard error of mean (SEM) are shown. A P-value significance threshold of < 0.05 was used, with levels of significance represented by *p < 0.05, **p < 0.01 ***p < 0.01 ****p < 0.001.

## Resource Availability

### Lead contact

Requests for further information and resources should be directed to and will be fulfilled by the lead contact, Richard Baines.

### Materials availability

This study did not generate new unique reagents. Fly stocks used are available on request from the lead contact without restriction.

### Data and code availability

Figures display all raw data values in addition to means ± SEM. All data reported in this report will be shared by the lead contact upon reasonable request.

This paper does not report original code.

Any additional information required to reanalysing the data reported in this report is available from the lead contact upon reasonable request.

### Funding

This research was funded through a joint Wellcome Trust investigator award (Grant 217099/Z/19/Z) between R.B, and M.L. Work on this project benefited from the Manchester Fly Facility, established through funds from the university and the Wellcome Trust (087742/Z/08/Z).The work also benefited from the University of Cambridge Imaging Facility, Department of Zoology, supported by Matt Wayland, and funds from a Wellcome Trust Equipment Grant (WT079204) with contributions by the Sir Isaac Newton Trust in Cambridge, including Research Grant [18.07ii(c)].

### Author contributions

Conceptualisation, B. C., J. D., & R. A. B.; methodology, B. C., J. D., R. B., & T. P.; Investigation, B. C., J. D., & T.P; writing – original draft, B. C.; writing – review & editing, B. C, R. A. B., & T. P., funding acquisition, R. A. B. & M. L., resources, R. A. B. & M. L.; supervision R. A. B. & M. L.

## Acknowledgements

We are grateful to the members of both the Baines and Landgraf labs for their guidance and collaboration. We thank Iain Hunter and Adam Bradlaugh for their feedback and constructive discussions throughout this work.

## STAR+METHODS

### Key Resources Table

#### Experimental Model and Study Participant Details

##### Method Details

- *Drosophila rearing and stocks*
- *Developmental activity manipulation*
- *Whole cell patch clamp electrophysiology*
- *Sample preparation for immunofluorescence*
- *Immunofluorescence staining of nerve cords*
- *Immunofluorescence imaging*
- *Quantification and Statistical Analysis*

---

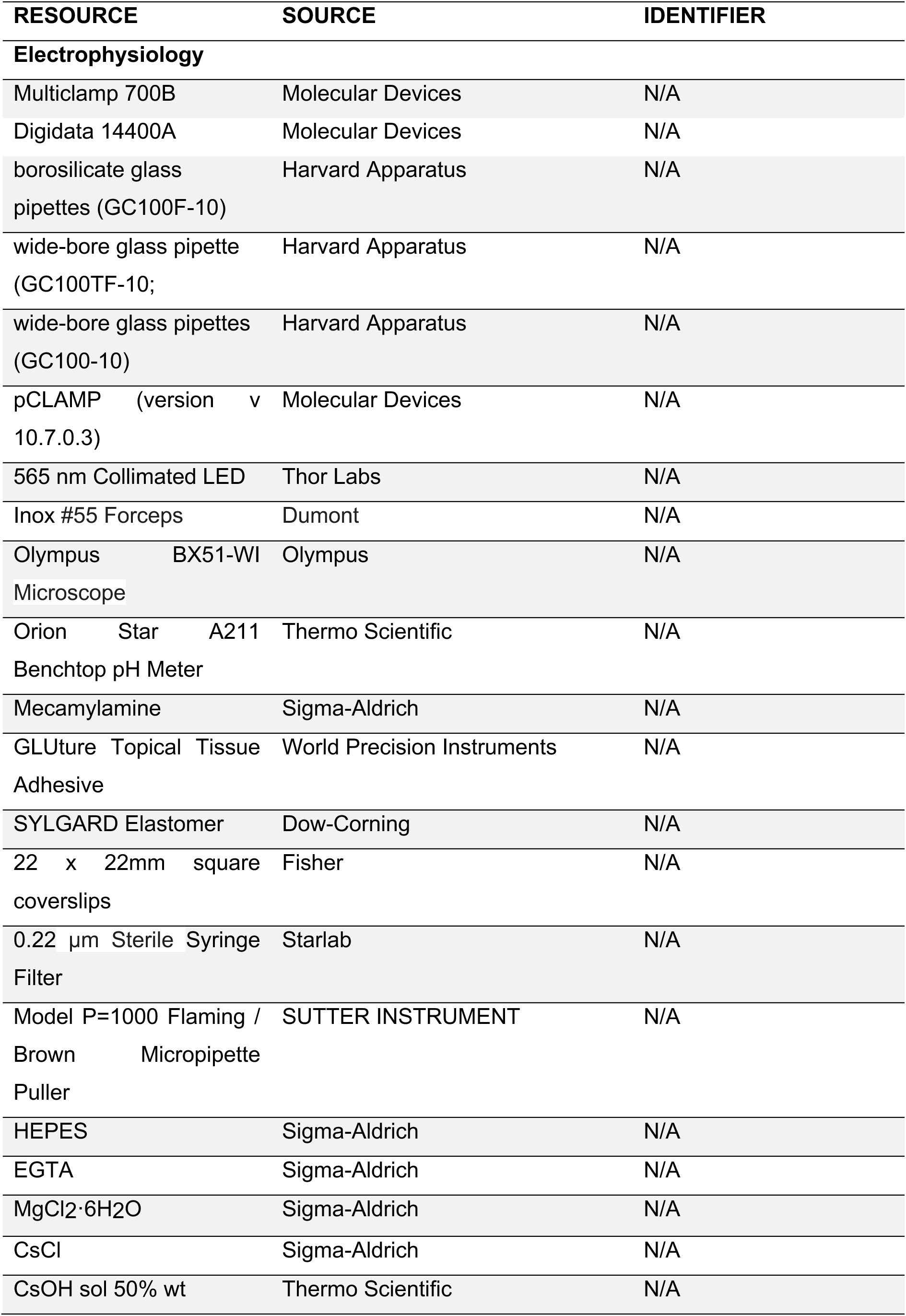
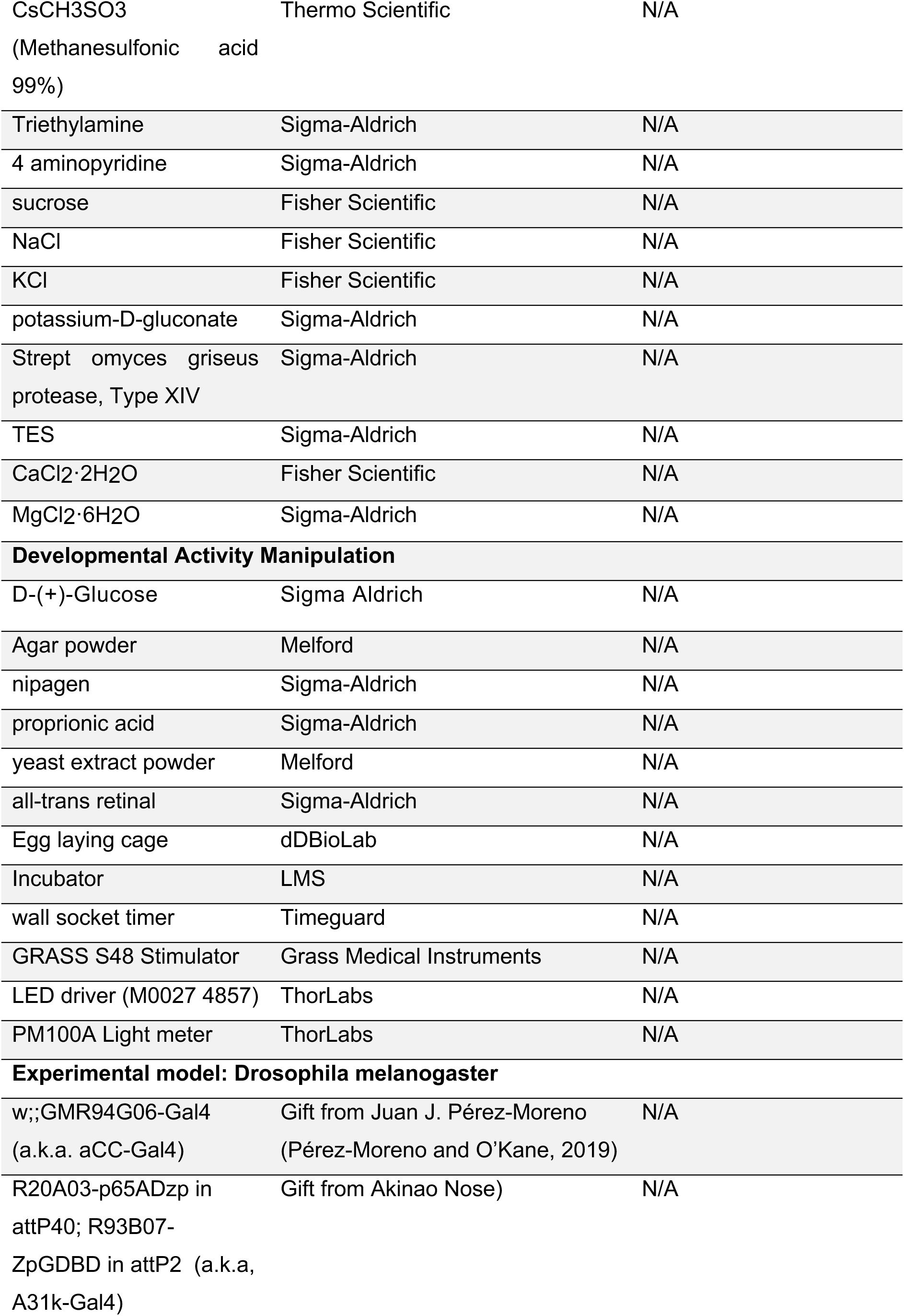
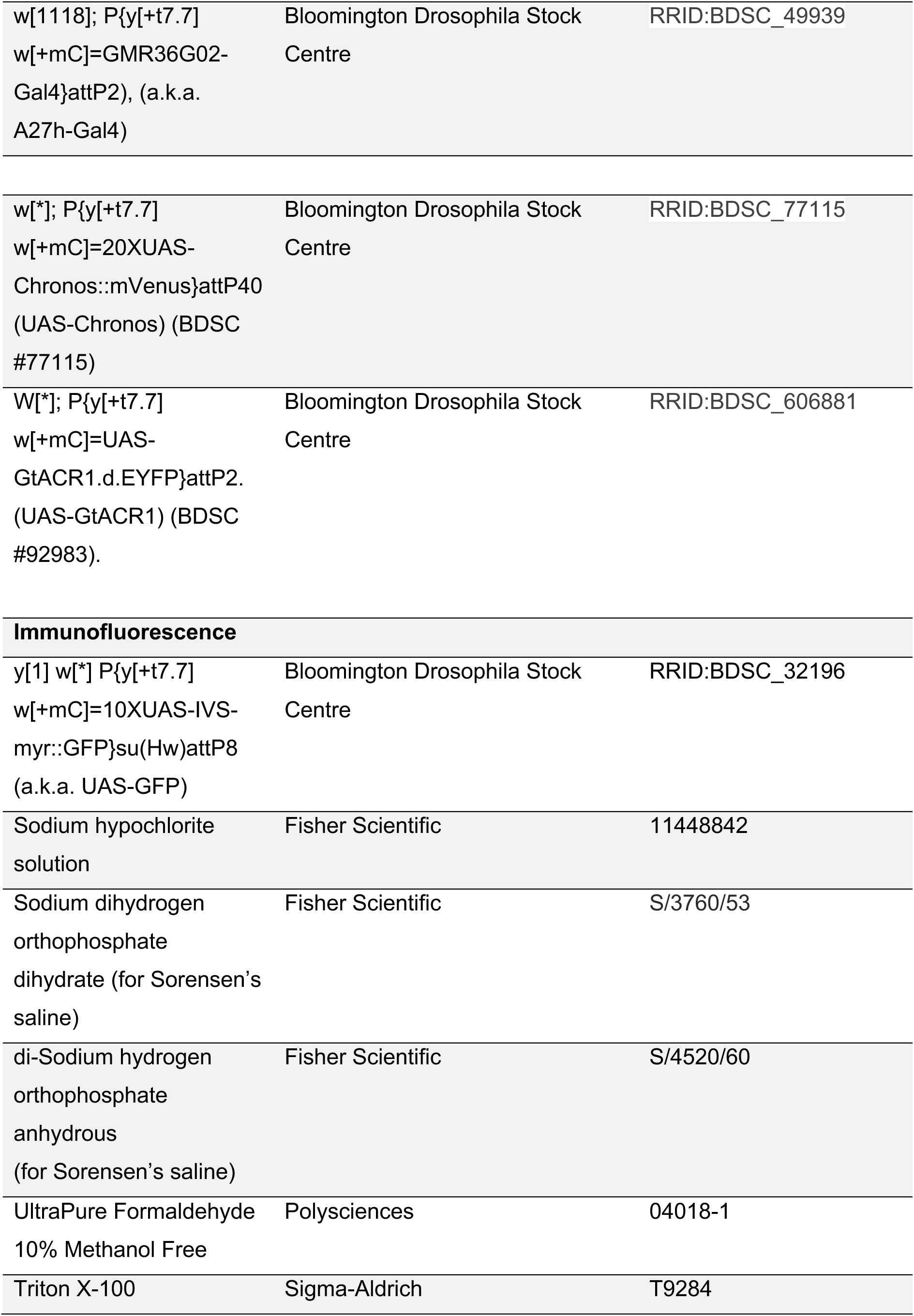
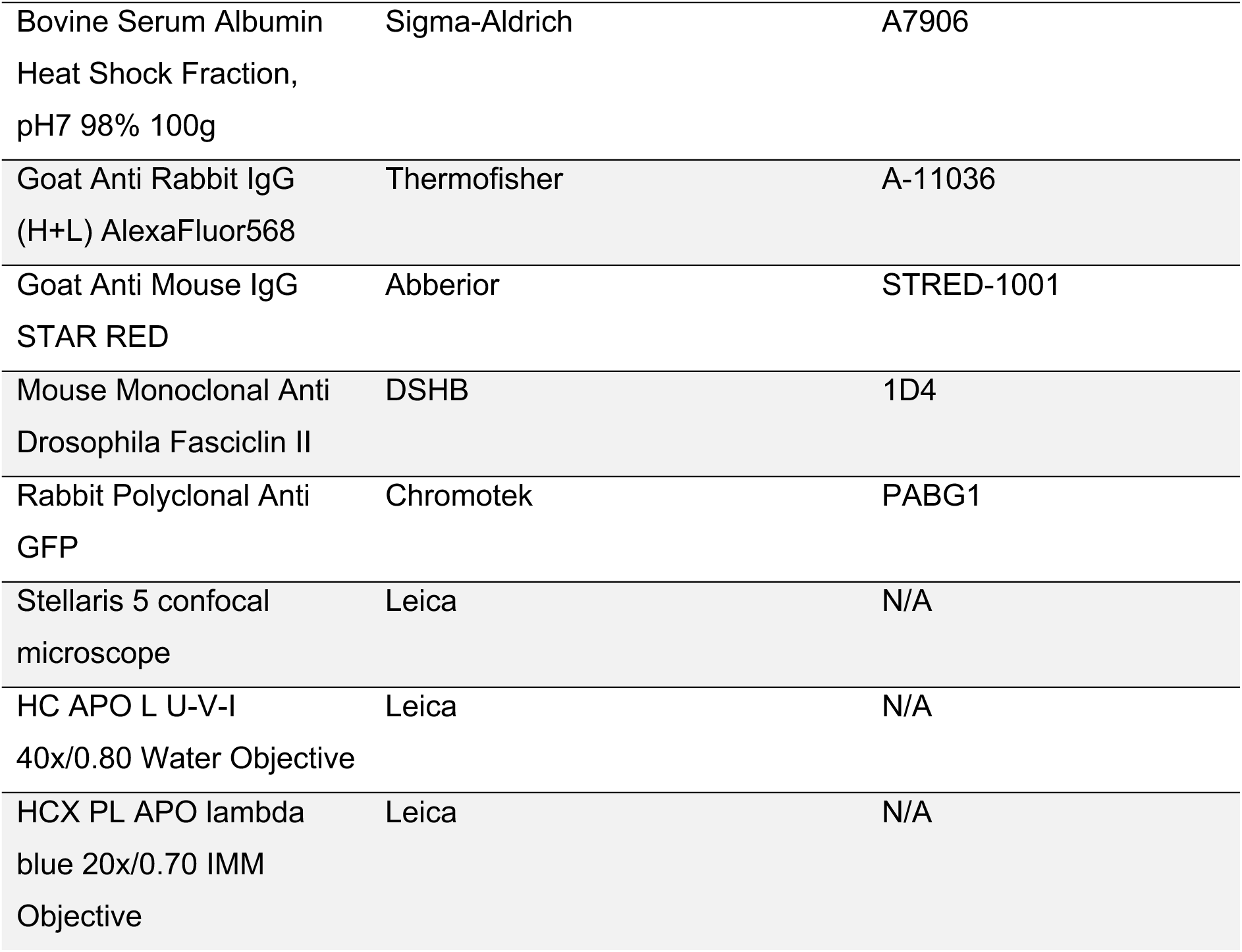

