## Supplementary for "Neuronal excitability is permanently altered by activity manipulation during an embryonic critical period in *Drosophila*"

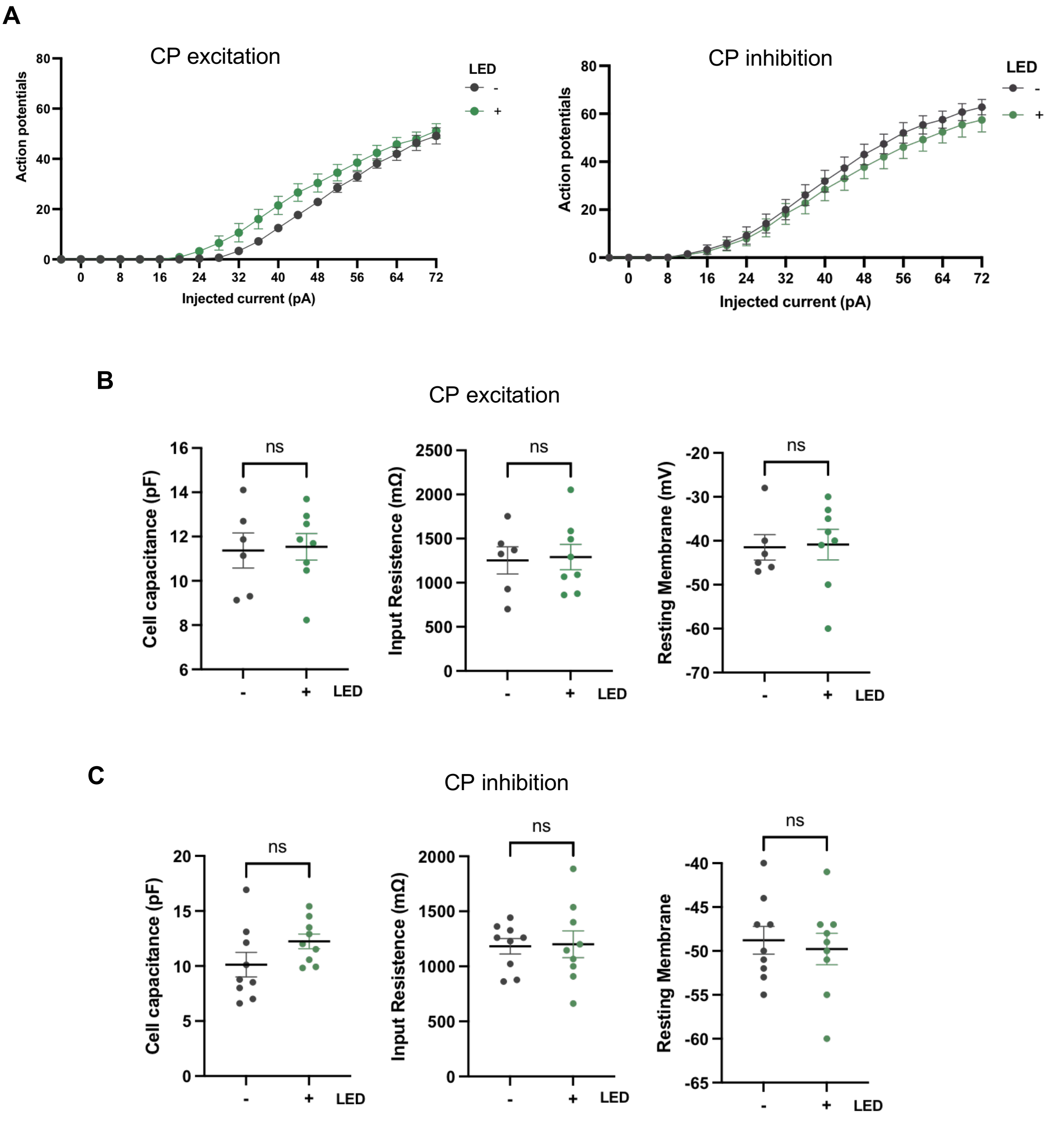


Supplementary 1: A27h excitability across the entire current step range used, with no change in cell capacitance, input resistance, or membrane potential upon break in.

**A**: A27h excitability across the entire current step range used for both non-manipulated controls (-LED) and larvae subjected to optogenetic excitation or inhibition (+LED) during the CP. **B**: No difference in A27h cell capacitance (left panel: unpaired t test, $t(12) = 0.168$, $p = 0.869$, $n = 6,8$), input resistance (centre panel: unpaired t test, $t(12) = 0.172$, $p = 0.867$ $n = 6,8$), or resting membrane potential upon break in (right panel: unpaired t test, $t(12) = 0.132$, $p = 0.897$, $n = 6,8$) when comparing between non-manipulated controls and larvae subjected to optogenetic excitation during the CP. **C**: No difference in A27h cell capacitance (left panel: unpaired t test, $t(16) = 1.279$, $p = 0.219$, $n = 9,9$), input resistance (centre panel: unpaired t test, $t(16) = 0.036$, $p = 0.972$, $n = 9,9$), or resting membrane potential upon break in (right panel: unpaired t test, $t(16) = 0.42$, $p = 0.68$, $n = 9,9$) when comparing between non-manipulated controls and larvae subjected to optogenetic inhibition during the CP.

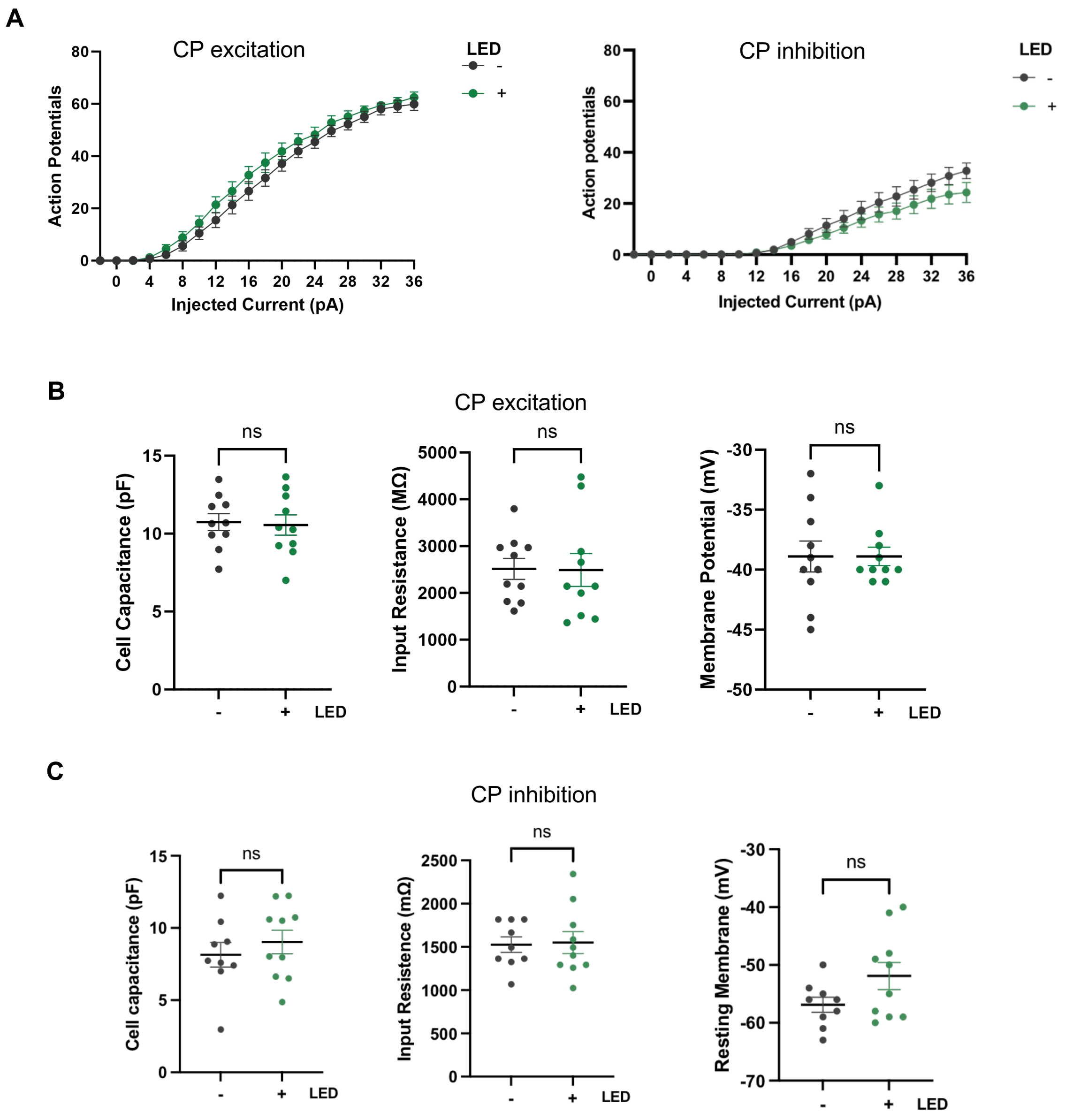


Supplementary 2: A31k excitability across entire current step range, with no change in cell capacitance, input resistance, or membrane potential upon break in.

**A**: A31k excitability across the entire current step range used for both non-manipulated controls (-LED) and larvae subjected to optogenetic excitation or inhibition (+LED) during the CP. **B**: No difference in A31k cell capacitance (left panel: Mann-Whitney test $U = 46$, $p = 0.796$, $n = 10,10$), input resistance (centre panel: unpaired t test, $t(17) = 0.058$, $p = 0.954$, $n = 10,10$), or resting membrane potential upon break in (right panel: unpaired t test, $t(17)$ <0.001, $p = 0.99$, $n = 10,10$) when comparing between non-manipulated controls and larvae subjected to optogenetic excitation during the CP. **C**: No difference in A31k cell capacitance (left panel: unpaired t test, $t(17) = 0.749$, $p = 0.464$, $n = 9,10$), input resistance (centre panel: unpaired t test, $t(17) = 0.153$, $p = 0.88$, $n = 9,10$), or resting membrane potential upon break in (right panel: unpaired t test, $t(17) = 1.795$, $p = 0.09$, $n = 9,10$) when comparing between non-manipulated controls and larvae subjected to optogenetic inhibition during the CP.

**A**

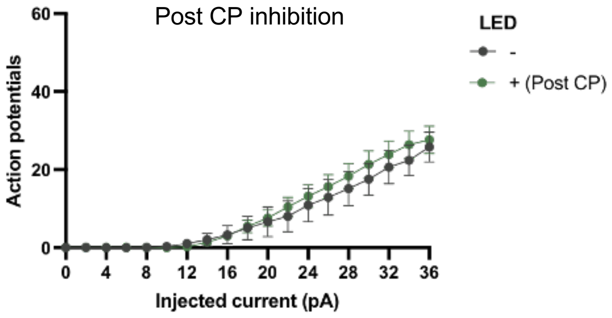


**B**

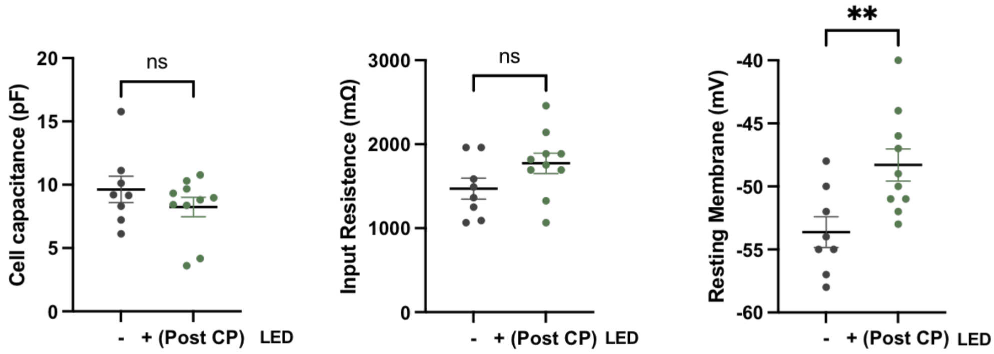


Supplementary 3: A31k excitability across the entire current step range used, cell capacitance, input resistance, and resting membrane potential on break in, comparing non-manipulated controls to post CP manipulation.

**A**: A31k excitability across the entire current step range used for both non-manipulated controls (-LED) and larvae subjected to optogenetic excitation (+LED) immediately after the CP closed at 19-21 h AEL. **B**: No difference in A31k cell capacitance (left panel: unpaired t test, $t(16) = 1.376$, $p = 0.185$, $n = 8,10$) or input resistance (centre panel: Mann-Whitney test $U = 24.5$, $p = 0.18$, $n = 8,10$) between conditions. Resting membrane potential upon break in (right panel: unpaired t test, $t(16) = 2.961$, $p = 0.009$, $n = 8,10$) was increased in larvae subjected to optogenetic excitation post CP compared to non-manipulated controls.

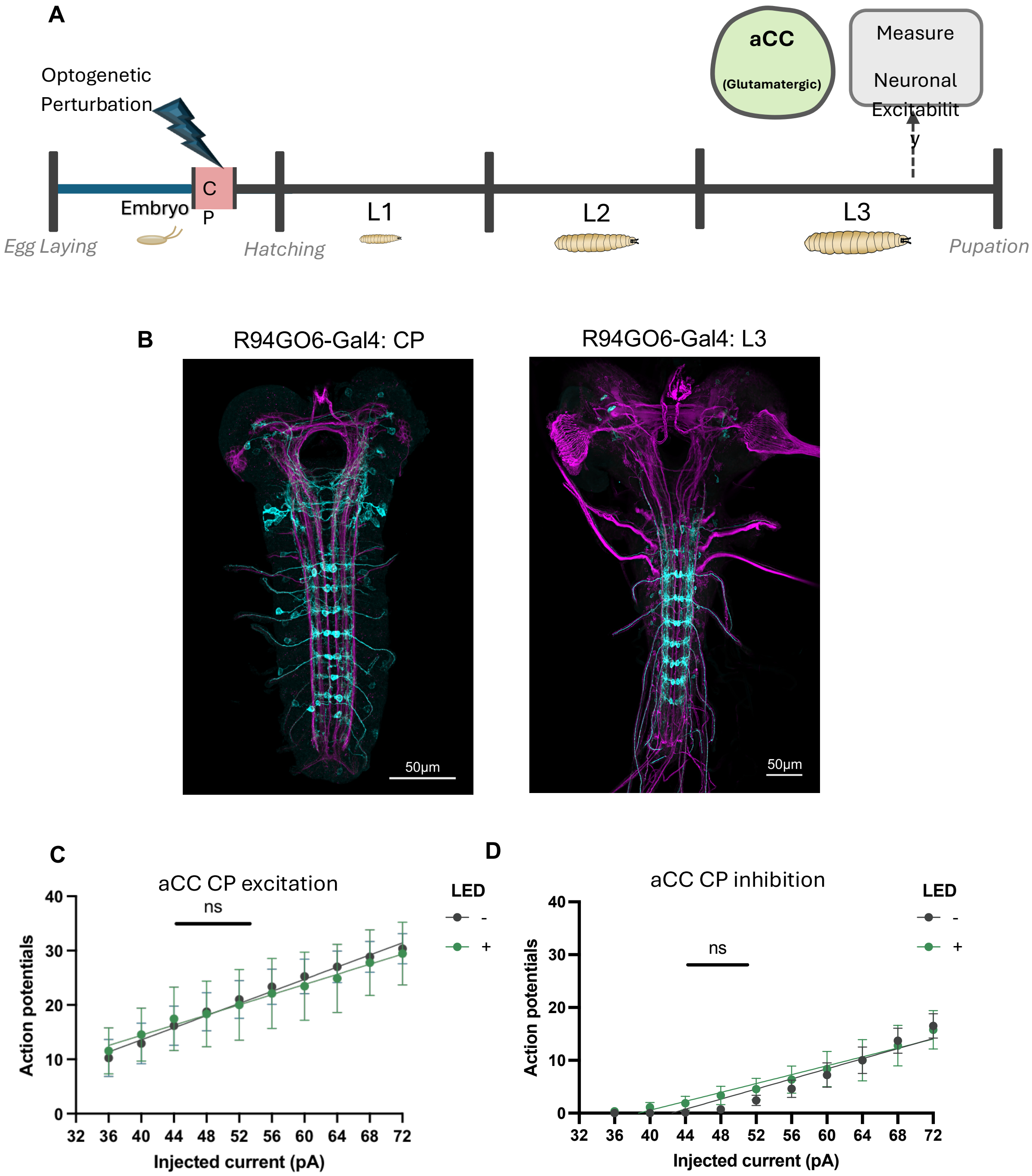


Supplementary 4: Optogenetic manipulation of aCC (MN1b) motoneuron between 17-19 h AEL does not alter intrinsic excitability measured at L3.

**A**: aCC motoneurons were optogenetically excited or inhibited during the embryonic critical period. aCC neurons were then patched at L3 to compare excitability to non-manipulated controls. **B**: Gal4/UAS expression in aCC motoneurons at 19 h AEL (left), and 72 h AEL (right) in the VNC. **C**: Optogenetic excitation between 17-19 h AEL does not alter intrinsic excitability of aCC motoneurons in L3 larva (-LED: slope 0.47, $R^2$ 0.09, n = 9, + LED: slope 0.56, $R^2$ 0.26, n = 12, comparison of intercepts, (F(1, 207) = 0.05, p = 0.824)). **D**: Optogenetic inhibition between 17-19 h AEL does not alter intrinsic excitability of aCC motoneurons in L3 larva (-LED: slope 0.48, $R^2$ 0.53, n = 10, + LED: slope 0.42, $R^2$ 0.29, n = 9, comparison of intercepts, (F(1, 187) = 0.96, p = 0.33)).

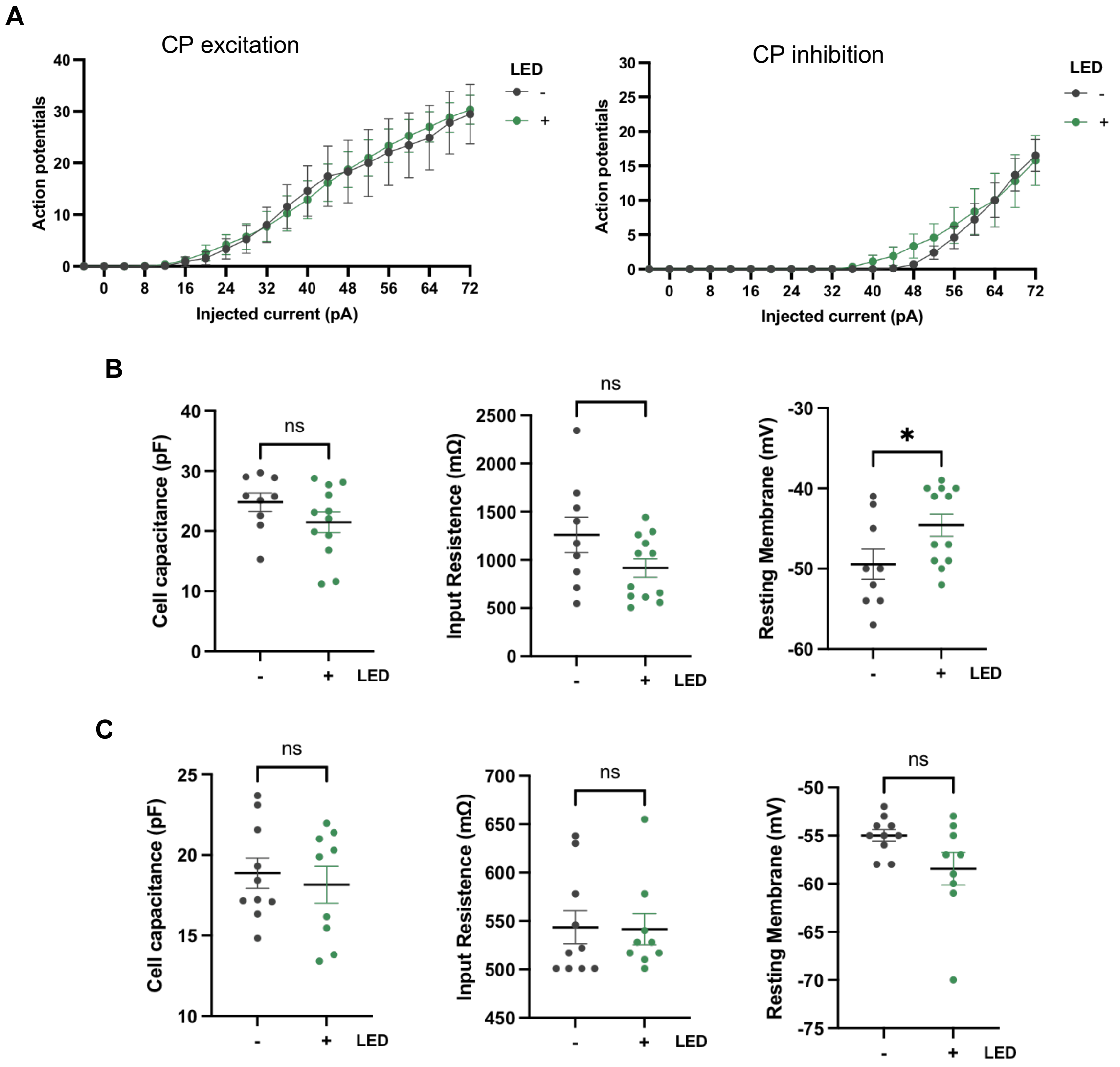


Supplementary 5: aCC excitability across the entire current step range used, with no change in either cell capacitance or input resistance but resting membrane potential upon break in being increased after CP excitation.

**A**: aCC excitability across the entire current step range used for both non-manipulated controls (-LED) and larvae subjected to optogenetic excitation or inhibition (+LED) during the CP. B: No difference in aCC cell capacitance (left panel: unpaired t test, $t(19) = 1.376$, $p = 0.185$, $n$ = 9,12) or input resistance (centre panel: unpaired t test, $t(19) = 1.767$, $p = 0.093$, $n$ = 9,12) when comparing between non-manipulated controls and larvae subjected to optogenetic excitation during the CP**.** Resting membrane potential upon break in was increased after CP optogenetic excitation (right panel: unpaired t test, $t(19) = 2.134$, $p = 0.046$, $n$ = 9,12)**. C**: No difference in aCC cell capacitance (left panel: unpaired t test, $t(17) = 0.49$, $p = 0.631$, $n$ = 10,9), input resistance (centre panel: Mann-Whitney test, $U = 39.5$, $p = 0.671$, $n$ = 10,9), or resting membrane potential upon break in (right panel: unpaired t test, $t(17) = 1.986$, $p = 0.063$, $n$ = 10,9) when comparing between non-manipulated controls and larvae subjected to optogenetic inhibition during the CP.

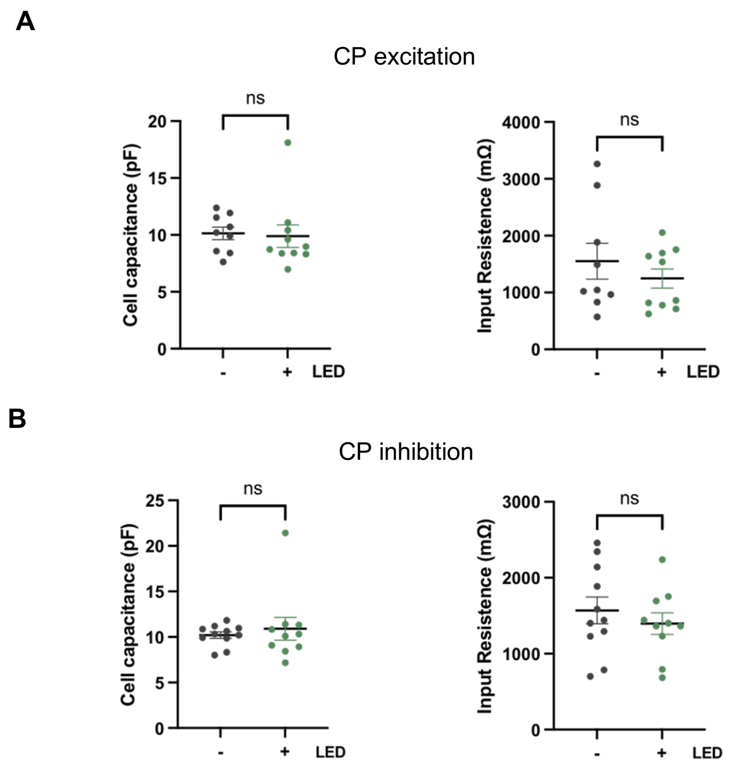


Supplementary 6: A31k sodium current recordings, no change in cell capacitance and input resistance between CP manipulated larva and non-manipulated controls.

**A**: No difference in A31k cell capacitance (left panel: Mann-Whitney test, $U = 32.5$, $p = 0.326$, $n = 9{,}10$), or input resistance (centre panel: unpaired t test, $t(17) = 0.872$, $p = 0.395$, $n = 9{,}10$) between non-manipulated controls and CP excitation. **B**: No difference in A31k cell capacitance (left panel: unpaired t test, $t(19) = 0.573$, $p = 0.573$, $n = 11{,}10$), or input resistance (centre panel: unpaired t test, $t(19) = 0.757$, $p = 0.459$, $n = 11{,}10$) between non-manipulated controls and CP excitation. Because these recordings were conducted in voltage clamp, membrane potential was not measured (but set to -60mV).
